# A high-throughput 3D cantilever array to model airway smooth muscle hypercontractility in asthma

**DOI:** 10.1101/2022.10.25.513767

**Authors:** Pranjali Beri, Christopher Plunkett, Joshua Barbara, Chien-Cheng Shih, S. Whitney Barnes, Olivia Ross, Paula Choconta, Ton Trinh, Bella Litvin, John Walker, Minhua Qiu, Scott Hammack, Erin Toyama

**Affiliations:** Novartis Institutes for Biomedical Research, San Diego, 92121, USA

**Author notes:** these authors contributed equally to this work (Pranjali Beri, Christopher Plunkett).

**Keywords:** asthma, smooth muscle, contractility, airway remodeling, cytokines

## Abstract

Asthma is often characterized by tissue-level mechanical phenotypes that include remodeling of the airway and an increase in airway tightening driven by the underlying smooth muscle. Existing therapies only provide symptom relief and do not improve the baseline narrowing of the airway or halt progression of the disease. To investigate such targeted therapeutics, there is a need for models that can recapitulate the 3D environment present in this tissue, provide phenotypic readouts of contractility, and be easily integrated into existing assay plate designs and laboratory automation used in drug discovery campaigns. To address this, we have developed DEFLCT, a high-throughput plate insert that can be paired with standard labware to easily generate high volumes of microscale tissues *in vitro* for screening applications. Using this platform, we exposed primary human airway smooth muscle cell-derived microtissues to a panel of six of inflammatory cytokines present in the asthmatic niche, identifying TGF-β1 and IL-13 as strong contractile modulators. RNAseq analysis further demonstrated enrichment of contractile and remodeling-relevant pathways in TGF-β1 and IL-13 treated tissues as well as pathways generally associated with asthma. Taken together, these data establish a disease relevant, 3D tissue model for the asthmatic airway which combines niche specific inflammatory cues and complex mechanical readouts that can be utilized in drug discovery efforts.

## Introduction

Chronic asthma is characterized by permanent structural changes to the airways known as airway remodeling, which consists of modifications to epithelial cells, smooth muscle, and fibroblasts as well as changes in the extracellular matrix (ECM) composition^1^. More recently, the focus of therapeutic drug discovery in asthma has shifted to the role of smooth muscle^2^, whose structural changes have been linked to hyperplasia, hypertrophy, and increases in bronchoconstriction^1,3^. The latter consists of both a baseline increase in airway tightness and narrowing due to smooth muscle hypercontractility^3–5^—which goes hand in hand with an increased resistance to airway relaxation in asthmatics compared to healthy patients—as well as increased sensitivity in response to triggers of bronchoconstriction^3^. In cases of severe asthma (5-10% of the patient population) with poor response to inhaled corticosteroids or bronchodilators such as LABA/LAMAs (long-acting beta-agonist/long-acting muscarinic antagonist)^6,7^, bronchial thermoplasty, or the removal of excess smooth muscle to improve baseline relaxation of the airway, is an option but this procedure is risky and might not provide long term relief^7^. Therefore, it is essential to identify therapies to reduce the aberrant shortening induced by hypercontractile airway smooth muscle present in asthma.

A major challenge in drug discovery when screening for complex mechanical phenotypes such as hypercontractility is the lack of high throughput systems that can both simulate the desired mechanics *in vitr*o and be easily integrated with existing automation systems and high-content imagers^5,8,9^. While there are several hydrogel-based *in vitro* models for modeling contractility such as simple gel contraction assays^10^ and traction force microscopy^11^, miniaturization is limited by design challenges such as poor attachment of the gel to the well bottom, interference of gel meniscus with imaging, complex analysis protocols, and low signal to noise ratios. There is also difficulty in scaling down to high-throughput arrays due to poor integration with existing high-content imagers and liquid handlers. Additionally, the current push for more physiologically relevant phenotypic screening requires the incorporation of complex 3D structures with human cells to increase the translatability of preclinical data and accelerate drug discovery^12,13^.

Elastomer-based cantilever strain gauges have emerged as a powerful means to model gross multi-cellular contractile forces in 3D at both the macro and microscales^14^. An improvement in the design of these tools is the inverted “hanging” style of construction whereby tissues are suspended from cantilevers that are mounted to the top of the cell culture multi-well plate which greatly simplifies tissue casting and improves imaging resolution^15,16^. These bioengineered tissue systems however are often lower in total throughput than highly multiplexed experiments require and utilize formats incompatible with existing microwell plate-based laboratory robotics and instrumentation^17^. The result is a lack of available 3D systems that can be easily integrated into existing screening protocols which in turn limits the scope of drug discovery-focused experimental designs and broad platform adoption for target ID studies.

To address these challenges, we have developed a 96-well <u>d</u>evice <u>e</u>nabling <u>f</u>unctional <u>l</u>inear <u>c</u>ontractility of <u>t</u>issues (DEFLCT) for multiplexed disease modeling of microtissue structures comprised of contractile cells. This system is simple to construct in high volumes, fully compatible with off the shelf assay plates, and leverages hanging construction for reproducible tissue casting. We then utilized this platform to expose airway smooth muscle cell microtissues to a panel of inflammatory cytokines to induce smooth muscle shortening *via* increased contractility. Lastly, we validated the shift towards an asthmatic disease phenotype through RNAseq, which revealed enrichment of pathways in the remodeled tissues that are associated with asthma, airway remodeling, and contractility.

## Results

### Development of a 96 Well ANSI/SLAS Compatible Cantilever System

To achieve sufficient assay throughput for parallelized multi-stimuli screening of contractility, we developed a 96-well format cantilever array to culture airway smooth muscle 3D microtissues. The device functions as a plate insert hanging down into the individual wells which allows for the use of standard ANSI/SLAS (Society of Biomolecular Screening) dimensioned labware and existing high content laboratory instruments (Figure 1A and Supplemental Figure 1A). To achieve this high density of microscale features while maintaining physiologically relevant rigidity, we elected to utilize NuSil MED-4940 silicone resin which has an equivalent stiffness to the more commonly utilized Dow-Corning Sylgard 184 but maintains a 3-fold higher ultimate tensile strain allowing for aggressive demolding without feature damage (Supplemental Figure 1B-D). Computational modeling of this cantilever design yielded an estimated spring constant of 0.109μN/μm and demonstrated symmetrical deflections in response to applied force (Figure 1B and C). Due to the significantly higher viscosity of MED-4940 precursor resin, pressurized molding into an aluminum form was employed to fabricate consumables (Supplemental Figure 2A).

**Figure 1:**
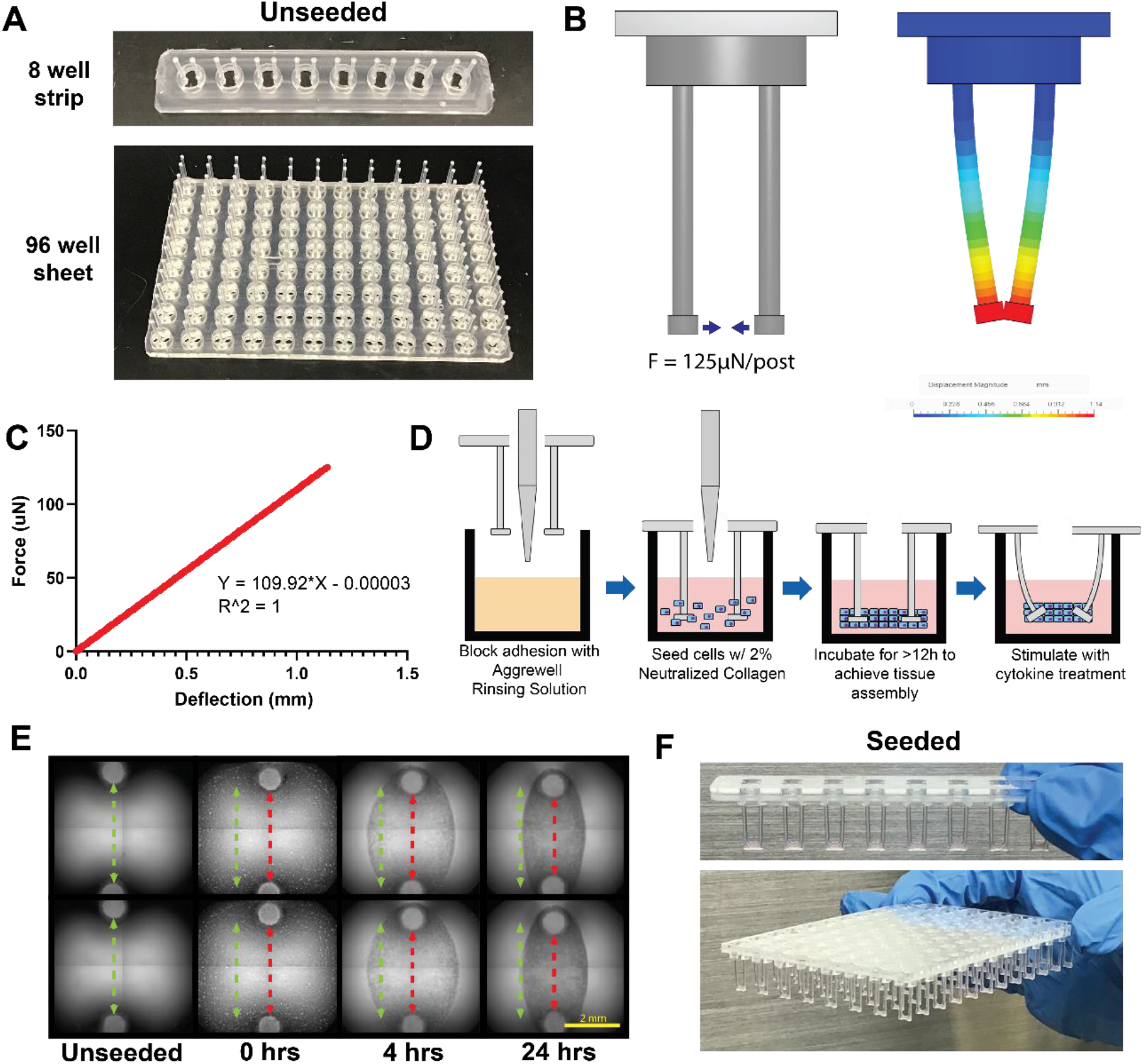
Overview of DEFLCT system design and general performance. **A)** Fully assembled DEFLCT consumables unseeded for 8 and 96 well formats. Parts are constructed from a silicone elastomer pillar array coupled to a rigid acrylic backing allowing for easy handling and placement inside standard ANSI/SLAS compliant assay plates. Individual 8 well column strips are utilized for limited throughput studies while full 96 well arrays are intended for screening experiments. **B)** Finite elements modeling of DEFLCT cantilever design bending in response to 125μN of applied force, the theoretical maximum readout of the sensor. **C)** Cantilever force-distance curve derived from FE simulations demonstrating a system spring constant of 109.92μN/mm per cantilever determined by linear regression. **D)** Process overview for seeding DEFLCT systems. Standard 96 well labware is coated with an adhesion blocking buffer after which cells are added to the well along with a pre-polymerized hydrogel. The gel is then allowed to cure around the posts forming the tissue and causes baseline deflection of the cantilevers within 24h. **E)** Timelapse image of tissue formation and tightening on cantilevers over a 24h period. Initial cantilever spacing (green) is reduced over the formation period during consolidation of the hydrogel and tightening of the seeded cells (red). **F)** Seeded DEFLCT consumables removed from their assay plates with visible microtissues spanning all cantilever pairs. Scale bar = 2mm.

Tissue casting on DEFLCT followed a straightforward four step process where, following adhesion blocking of the assay plate, a mixture of cells and neutralized collagen was added to the wells. Then, the DEFLCT consumable was inserted into the plate and the mixture was allowed to polymerize for 15 minutes at 37°C before adding growth media and incubating overnight (Figure 1D). Multiple seeding conditions were tested to optimize microtissue formation, with the tightest tissues derived from 50,000 cells suspended in a 1mg/mL collagen gel solution (Supplemental Figure 3). Given these conditions, tissues began compacting within 4h of initial seeding and had fully consolidated by 24h (Figure 1E). Applied across a full assay plate, this casting method was able to achieve successful tissue formation on all 96 cantilever pairs across multiple patient derived doner cell lines (Supplemental Figure 4). Consumables were fabricated in 8-well partial plate strips and 96-well full-plate formats to allow for both small- and large-scale experiments (Figure 1F).

### TGF-β1 and IL-13 Induce Contractile Phenotypes in Patient Derived hBSMCs

The compatibility of this system with existing well-plates and the 96-well format enables high-throughput, parallelized investigation of the effects of chemical stimuli (compounds, chemokines, etc.) on 3D microtissues of contractile cells to generate disease models and investigate potential therapies for diseases involving contractile cells. We applied these high-throughput capabilities to study changes in the contractile phenotype of primary human bronchial smooth muscle cells after treatment with TGF-β1^18^, IL-13^19^, TNFα^20^, IL-1β^21^, IL-17A^8^, and INFγ^22^, a panel of six inflammatory cytokines associated with airway inflammation in asthma and linked to airway smooth muscle contractility in previous literature.

Microtissues of primary human bronchial smooth muscle cells (hBSMCs) from 4 independent healthy donors were seeded in a 96 well plate with the DEFLCT full plate insert, and tissues were observed in all wells after 24 hours (Figure 2A and Supplemental Figure 4). Tissues with DEFLCT were manually transferred to a new plate containing starvation media (0.5% FBS) and the panel of inflammatory cytokines (all at 10ng/mL) (Figure 2A). Microtissues were cultured until significant tissue shortening was observed in the treatment conditions, which was apparent by Day 7. Thus a 7-day timeframe was used as the experimental endpoint (Figure 2A and Supplemental Figure 5). A trained deep learning model was used to measure the Day 7 distance between the cantilever tips (T1) and earlier, the unseeded Day 0 distance (T0) (Figure 2B). The difference in distances was converted to applied force using the equation in Figure 1C.

**Figure 2:**
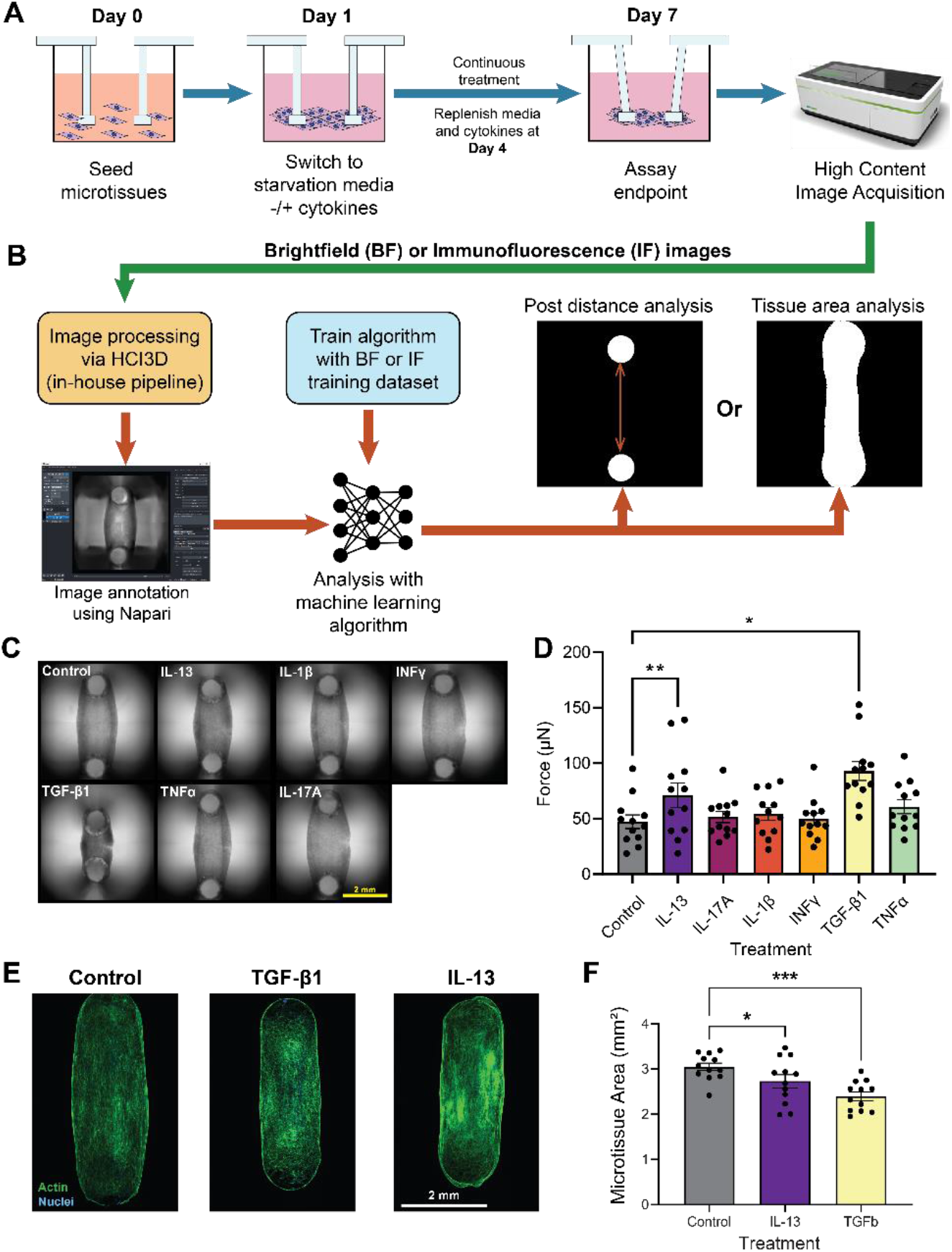
TGF-β1 and IL-13 induce contractile phenotypes in bSMC microtissues cultured on DEFLCT arrays. **A)** Graphical abstract of experimental procedure and key assay timepoints. **B)** Overview of machine vision image processing approach for contraction and tissue area readout. **C)** Representative brightfield images of tissues treated with experimental cytokines at day 7. Scale Bar = 2mm. **D)** Contractile forces exerted by microtissues for cytokine treatment conditions. All individual data points represent biological replicates that were averaged from ≥ 3 technical replicates each. Conditions are comprised of 3 biological replicates for 4 human donors (n=12 microtissues per condition). Data represented as mean ± S.E.M. and analyzed by 1 way ANOVA with Geisser-Greenhouse correction and Dunnet’s multiple comparisons. *p<0.05; **p<0.01. **E)** Representative F-actin (green) and nuclear (blue) staining of cytokine conditions associated with increased contractility and control group. Scale bar = 2mm. **F)** Total microtissue area as measured by F-Actin fluorescence thresholding. All individual data points represent biological replicates. Conditions are comprised of 3 biological replicates for 4 human donors (n=12 data points per condition). Data represented as mean ± S.E.M. and analyzed by 1 way ANOVA with Geisser-Greenhouse correction and Dunnet’s multiple comparisons. *p<0.05; ***p<0.001.

We observed that hBSMCs treated with TGF-β1 and IL-13 showed, on average, the greatest decreases in distance between the cantilevers and applied significantly higher contractile forces to the cantilevers compared to the healthy controls (Figure 2C and D). Fluorescent staining of the tissues and measurement of the tissue area after fixing and staining for F-actin confirmed that the total tissue area decreases in hBSMC microtissues treated with TGF-β1 and IL-13 compared to control tissues, indicating that the tissue is tightening with treatment (Figure 2E and F). These data combined indicate that we were able to drive microtissues of healthy cells towards a hypercontractile shortened state that is present in asthmatic airways, thus identifying the asthma-associated inflammatory cues that could generate a diseased microtissue phenotype whose contractility can be monitored on this system.

### TGF-β1 and IL-13 cause a shift in the transcriptomic contractility signature

To investigate the transcriptomic changes that are underlying the increased contractility and decreased tissue area observed after TGF-β1 and IL-13 treatment, we performed bulk RNA sequencing on microtissues from 4 independent healthy donors that were treated with 10ng/mL of TGF-β1 and IL-13. While both TGF-β1 and IL-13 caused significant increases in applied force and decreases in tissue area, they differed drastically in their differentially expressed genes (DEGs) signature. TGF-β1 treatment resulted in 3688 DEGs and IL-13 treatment resulted in 138 DEGs that had a p-value less than 0.05 and a fold change greater than 1.5 (|Log2FC| > 0.585) (Figure 3A and B, Supplemental Tables S2-4). DEGs that were upregulated in each condition were input into Enrichr^23^ to obtain the key pathways of interest. Alignment to the KEGG pathways database and the human MSigDB Hallmark collections revealed several overlapping pathways of interest in both TGF-β1 and IL-13 treated microtissues (Figure C and D). The full list of pathways can be seen in Supplementary Tables S5-8.

**Figure 3:**
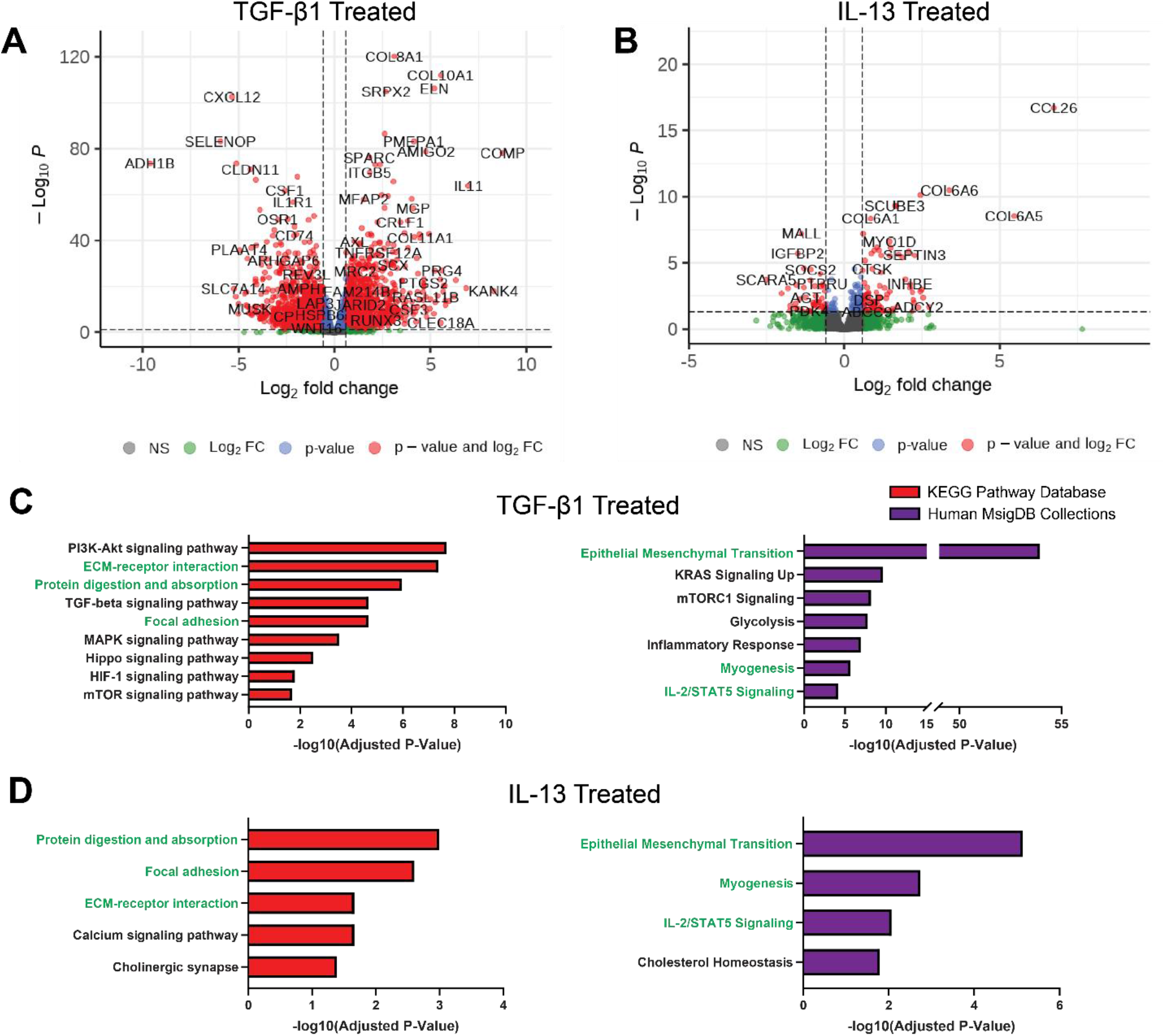
Contractility inducing cytokines independently enrich common pathways associated with asthmatic phenotypes. **A-B)** Bulk RNA sequencing volcano plots for differentially expressed genes in TGF-β1 **(A)** and IL-13 **(B)** treated microtissues. DEG’s are defined as having a p value >0.05 and a Log_2_ fold change > 0.585 (FC > 1.5). **C-D)** Select pathways enriched by DEG’s in TGF-β1 **(C)** and IL-13 **(D)** treated microtissues associated with an asthmatic SMC phenotype. All shown pathways were curated from the 20 lowest adjusted p-value results (with an absolute minimum adjusted p-value<0.05) from Enrichr pathway analysis using KEGG Pathway Database (red) and Human MsigDB Collections (purple) for DEG’s. Common pathways enriched in both TGF-β1 and IL-13 treated tissues are highlighted in green. A full list of all enriched pathways for each condition and pathway database along with specific associated DEG’s can be found in Supplemental Tables 2-8.

Both treatments resulted in an enrichment of pathways associated with protein digestion and absorption, focal adhesion, and extracellular matrix (ECM)-receptor interaction, which correlates with the observed phenotypic changes in the microtissues by the smooth muscle (Figure 3C and D). Closer observation of the top DEGs in both conditions show an upregulation in the expression of matrix proteins, including a variety of collagen subfamily proteins and elastin (ELN), as well as proteins involved in matrix modifications such as matrix metalloprotease 2 (MMP2) and lysyl oxidase (LOX), which can crosslink collagen fibers^24^ and is upregulated in both conditions (Supplementary Figure 6). The two treatment conditions also show an enrichment in epithelial to mesenchymal transition (EMT), the myogenesis pathway, and IL-2/STAT5 signaling (Figure 3C and D). (Figure 3C and D). Individually, TGF-β1 treatment shows an enrichment in PI3K-Akt, MAPK, Hippo, HIF-1, mTOR, and KRAS pathways, while IL-13 treated microtissues show a significant enrichment in calcium signaling, cholinergic synapse, and cholesterol homeostasis pathways. Cholesterol homeostasis is also significantly enriched in the TGF-β1 treated microtissue, which was not included in Figure 3 but can be seen in Supplemental Table S6. Combined with the observed phenotypic changes, these findings show pathways-level changes that might be responsible for the increased contractility. Furthermore, this model can be used for therapeutic drug discovery to treat shortened airways due to hypercontractile smooth muscle cells.

## Discussion

The narrowing of the airway by hypercontractile smooth muscle cells is a key feature of airway remodeling that distinguishes asthmatic and healthy airways^1,3,25,26^. To identify therapies to slow or halt the progression of disease or potentially aid in repair, it is essential to utilize a high-throughput system that can be used in phenotypic screening. We have developed a high-throughput platform for multiplexed 3D modeling of contractile microtissues of hBSMCs and applied this technology to investigate the impact of a panel of six inflammatory cytokines associated with asthma on hBSMC contractility. We found that only the TGF-β1 and IL-13-treated microtissues exhibited a significant increase in contractility, shown by an increase in applied force and resultant decrease in distance between the cantilevers, as well as a significant reduction in tissue area surrounding the posts. While all the tested cytokines included in the panel have been linked to modulation of airway smooth muscle contractility^8,18–22^, they have not previously been investigated as a panel with a single readout, which enables a cleaner determination of which individual cytokines are the strongest drivers of contractility and shortening in the airway smooth muscle. However, this system could be further applied to investigate synergistic effects of combinatorial cytokine treatments.

Previous studies of hypercontractile airway behavior have investigated changes in calcium flux or examined differences after contractile agonist treatment (i.e. histamine, acetylcholine), both of which provide information on acute exacerbation differences but have not captured baseline changes in smooth muscle contractility that would be consistent with airway remodeling^27,28^. Alternatively, baseline changes in the airway are often investigated with *in vivo* models or *ex vivo* slices, which are not practical for drug discovery biology^27–29^. The DEFLCT system was able to identify the important cytokine drivers of smooth muscle contractility and simultaneously provide an avenue to induce a disease phenotype for future investigations on therapeutics for repair without the need for tissue explants or expensive and time-consuming *in vivo* studies^27,29^. This technology could also potentially reduce the number of animals needed for drug discovery. Furthermore, its simple, robust fabrication process and adaptability to existing ANSI/SLAS labware and automation systems enables easy scalability to larger small molecule, siRNA, and CRISPR screening libraries in future studies.

In addition to inducing hypercontractility and microtissue shortening to phenotypically drive the hBSMCs towards an asthmatic airway phenotype, we employed RNAseq to investigate the transcriptomic changes that underly the shift in contractility. Pathways that were uniquely enriched in TGF-β1 treated microtissues included PI3K-Akt^30^, MAPK^31^, Hippo^32^, HIF-1^33^, mTOR^34^, and KRAS^35^, which have all been implicated in asthma. IL-13 treated microtissues also showed an enrichment in calcium signaling^36^, which plays an integral role in smooth muscle contraction. We found several overlapping pathways in the TGF-β1 and IL-13-treated microtissues which could explain the similarities in phenotypic changes. In both treatment conditions, upregulation in gene expression of ECM components and modifiers, as well as enrichment of pathways involved in focal adhesions and cell-ECM interactions supports the argument that the smooth muscle cells under the influence of these cytokines would play a hand in remodeling the airway matrix^24,37^. EMT has been previously implicated in asthma^38,39^ though not particularly in the context of airway smooth muscle given its already mesenchymal lineage^40^. However, upregulation in EMT-associated proteins could potentially signify a transition of the airway smooth muscle cells into a more contractile phenotype and further support a shift towards airway remodeling. Similarly, increases in pathways involved in myogenesis^41^, cholesterol homeostasis^42^, and STAT5 signaling^43^ can also be indicative of a shift towards the asthma phenotype, as all three pathways have been shown to play a role in allergic asthma inflammation and smooth muscle cell contractility. Given the link between many of the highlighted pathways with asthma and the changes in the contractile phenotype of the microtissues, we believe our system provides an important high-throughput avenue for investigation of therapeutics for the repair and reversal of airway smooth muscle hypercontractility and remodeling. The use to this technology can extend beyond upper airway respiratory diseases to any organ system where contractile cells play a role in or are affected by disease progression, including diseases involving the cardiovascular system^44^, fibrosis^45^, or the musculoskeletal system^46^.

There are technical limitations to the DEFLCT airway model which will require continued development of the system. The organization of *in vivo* smooth muscle tissue is radial rather than linear^3^, which is not directly recapitulated in the 3D hBSMC constructs and may limit translatability of exact force readouts. Additionally, this study does not address the role of crosstalk between the smooth muscle and other cells within the niche such as bronchial epithelial or T-helper cells. The easy transfer of DEFLCT consumables between tissue culture treated assay plates may enable future co-culture studies whereby smooth muscle constructs are suspended directly above monolayer cultures of disease relevant cell types. Finally, although the DEFLCT platform enables larger screening studies on the order of hundreds of well replicates, the use of larger and more diverse small molecule libraries common to pharma pipelines containing thousands or tens of thousands of compounds requires still higher throughput assay plate formats. Development of 3D contractility assays compatible with 384 and 1536 well formats could enable such screening experiments and MEMS technologies may offer a solution to the form factor challenges inherent to such a small volume.

## Conclusions

We have developed a high-throughput platform for multiplexed 3D microtissues that can provide quantitative measurements of contractility and be easily integrated with existing high-throughput screening platforms. We have applied this technology to understanding the effects of a panel of inflammatory cytokines on airway smooth muscle contractility and found that TGF-β1 and IL-13 induce a hypercontracted microtissue state compared to control. RNAseq analysis reveals that a majority of the distinct and overlapping enriched pathways across the two treatment conditions can be linked to asthma, smooth muscle contractility, and/or airway remodeling. Integration of complex, high-throughput 3D systems into investigating disease biology can help improve investigation of potential pathways for therapeutic intervention.

## Methods

### Cell Culture

Normal human bronchial smooth muscle cells (hBSMCs) were obtained from Lonza (CC-2576) and cultured using SmGM™-2 BulletKit™ (Lonza CC-3182). All cells were from patients ranging in age from 30 to 65 years old and were sub-cultured according to Lonza’s recommended protocol.

### Device Fabrication

The cantilever consumables were fabricated in either an 8w strip designed to fit into a microplate’s column or a full 96w array. The methods described represent fabrication techniques for both consumable sizes. Cantilever consumables were made by curing liquid silicone rubber (LSR) inside a machined aluminum mold. Equal amounts (1:1 mix ratio, parts A and B) of LSR (NuSil MED-4940) were mixed inside a vacuum planetary mixer (THINKY MIXER ARV-310P) at 1800rpm for a total approximate duration of 6 minutes. Once thoroughly mixed and degassed, a pneumatic dispensing gun (Semco® 250-A) was used to inject the LSR into a mold. The uncured LSR was vulcanized at 150°C for 2 minutes using a hydraulic lab press equipped with heated platens (Carver Bench Top Auto Press) then demolded from the tool.

A rigid acrylic backing sheet was attached to the sheet of soft silicone cantilevers to provide a stiff supportive layer facilitating ease of use. A tabletop laser cutter (Glowforge Pro) was used to cut the acrylic backing sheets for both types of cantilever consumables. A double-sided pressure sensitive adhesive (PSA) (3M 96042) was used to attach the silicone cantilevers to the laser cut acrylic backing sheets. The PSA was cut to the appropriate geometry using a craft cutter (Silhouette Cameo 4).

### Mechanical Testing and Finite Elements Modeling

Tensile testing samples were prepared as described above in 5mm × 24mm × 1mm rectangular strips. Samples were clamped into an Instron universal testing system (Instron, Norwood, MA) with a 500N load cell equipped. Samples were elongated at a rate of 5mm/min until the component failed by tearing. Young’s modulus and ultimate tensile strain were determined for each curve using Bluehill Universal software (Instron, Norwood MA).

FE model of the hanging posts was created in SimScale CAE software (SimScale GmbH). A second order finite mesh with 24,777 tetrahedral elements and 42,203 nodes was generated through an automatic mesh generator tool that is part of SimScale. Post material was considered to be incompressible, having Young’s modulus E = 1.33 MPa, Poisson’s ratio ν = 0.499 and density ρ = 1120 kg/m2. An inward force acting at the tip of the post was applied to induce deflection in the posts and simulate tissue contraction. Forces were symmetrically applied to each post to pull them together. Simulated deflection as a function of applied force was computed and plotted.

### Device Seeding and Cytokine Treatment

Cantilever arrays were initially treated with 100% oxygen plasma for 3min (Nordson Electronic Solutions, Concord, CA) then immediately submerged and incubated in a 0.15mg/mL type 1 collagen solution (Advanced Biomatrix 5153) for 1 hour to coat the silicone exterior. 96 well plates (Perkin Elmer 6055302) were pretreated with an anti-adhesion rinse solution (Stem Cell Technologies 07010) for a minimum of 15min to prevent cell attachment. Cells were then suspended in a 1 mg/mL neutralized type 1 collagen solution then dispensed at a density of 50,000 cells per well into the plate over ice. Cantilever arrays were immediately added to the well ensuring the ends of each cantilever were fully submerged and incubated at 37°C for 15 min to allow for gelation. Growth media was then added to the seeded wells and plates were incubated overnight to allow for initial tissue formation (Day 0).

Following 24h of incubation (Day 1), cantilever arrays with seeded microtissues were transferred into a fresh 96 well plate containing hBSMC growth media with reduced serum (0.5% FBS) and the indicated recombinant cytokines at 10 ng/mL (see Supplementary Table S1 for catalog numbers). Microtissues were incubated for 3 days, after which the media and cytokines were replenished (Day 4). After an additional three days of treatment (Day 7), microtissues were used for desired endpoint assessment.

### Contractile Force Analysis

Prior to seeding, brightfield images of the unseeded cantilever arrays were acquired on Day 0 using an Opera Phenix High-Content Imager (Perkin Elmer) to calculate the initial distance between the pillar caps (D0). At Day 7 of cytokine treatment, brightfield images of the cantilever arrays with tissues were acquired again to calculate the distance between the pillars after treatment (D7). After the Phenix acquired and exported images to storage servers, an in-house pipeline with the following functions was applied sequentially: background and shading correct (BaSiC)^47^, image stitching, and maximum intensity projection. The final outputs of 2160 × 2160 pixels were used during the evaluation of immunofluorescent staining. For cantilever post distance measurement, the image size was reduced to 540 × 540 pixels by 4 × 4 binning. The pipeline recorded pixel and voxel size (μm) from metadata for the downstream calculation. We performed data annotation in Napari^48^ by using a random-forest-based segmentation plugin named napari-accelerated-pixel-and-object-classification (APOC)^49^, paired with our customized image sampler and label cleaner, to create masks for both post and microtissue labeling.

Our analysis method integrates Medical Open Network for Artificial Intelligence (MONAI)^50^ and PyTorch^51^ for neural network training, utilizing the MONAI transform function to perform on-the-fly data augmentation in the training process. Selected transform functions include spatial crops (RandSpatialCropSamplesd), random flip (RandFlipd), scale and shift intensity (RandScaleIntensityd and RandShiftIntensityd), and 90 degrees rotate (RandRotate90d). Random probabilities for each transform function are assigned individually. We selected a cropping size of 192 × 192 pixels, sampling 40 times per image. In total, 173 datasets were used during model training.

The neural network architecture used in our method is two residual units, five layers of Res-UNet^52^. The loss function is the sum of Dice loss per batch in a batch size of 32. We performed our model training using an Nvidia Tesla V100-SXM2 GPU with 32 GB memory (Nvidia, Santa Clara, California). The training process ran within 500 epochs and was optimized by Adam optimizer^53^, with an initial learning rate of 0.001. 20% of training datasets were randomly chosen for validation, and our workflow selected the model with the highest Dice metrics for the following image segmentation.

Once the U-Net model predicted the regions of posts, we used rule-based methods, assisted by human inspection, to ensure the quality of segmentation results. Later, our approach generated the distance map with Euclidean distance transformation in SciPy^54^, detecting the minimum distance between two posts. The results are reported in both pixel and μm. Displacement was converted to mm, divided by 2 to get the displacement of one cantilever, and input into the calculated equation (Figure 1C) to obtain applied force per cantilever.

### Actin Staining and Analysis

Microtissues were transferred to wells with 4% PFA in PBS (diluted from 32% PFA solution, Electron Microscopy Sciences 15714-S) for 15 minutes. Tissues were then washed three times with PBS and blocked with 1% BSA (Millipore Sigma A2153) for 20 minutes, after which they were transferred to a staining solution containing Alexa Fluor 488 phalloidin (Thermo Fisher A12379) and Hoechst (Thermo Fisher 62249) and incubated overnight at room temperature on a shaker. Tissues were then washed three times with PBS and imaged at 5x magnification using an Opera Phenix High-Content Imager (Perkin Elmer) with four fields of view to capture the full microtissues, which were stitched before calculating total tissue area.

To calculate the area of the stained tissue, we employed a similar workflow to train the prediction model but with some modifications. Changes included: 1) Input images used for Res-UNet training were cropped from images with original pixel size and cropped into size of 384 × 384 pixels. 2) We scaled the intensity with a fixed range (ScaleIntensityRanged) calculated from our datasets 3) The training was run with a batch size of 6 within 600 epochs. With model predictions, our method applied region property measurement from scikit-image^55^ and returned area in pixel^2^ and mm^2^.

### RNA Sequencing

RNA was isolated from hBSMC microtissues that were either untreated or treated with TGF-β1 and IL-13 for 7 days prior to RNA isolation using the RNeasy Mini Kit (Qiagen 74106). The RNA yield was measured using Agilent TapeStation (4200 TapeStation System) at 260 nm. 200 ng of total RNA was used to make RNA sequencing libraries using the TruSeq Stranded mRNA Prep Kit (Illumina 20020595). Libraries were run on a 51-cycle NovaSeq (SP flowcell, 2 lanes) sequencing run and yielded an average sequencing depth of 8 million mapped reads per sample. Reads were generated from a NovaSeq instrument with bcl2fastq from Illumina, v 2.20.0.422-2 and were aligned to a genome and annotation from ensembl, assembly GRCh38.p13, release version 104^56^. Gene counts were assessed with RSEM v1.3^57^. Differential gene expression was assessed with DESeq2 v.1.28.1, in R version 4.2.0^58^. All conditions consist of RNA from four independent donors from n = 2-3 biological replicates. DEGs that were upregulated in each condition were input into Enrichr^23^ to obtain the key pathways of interest.

### Statistical Analysis

All experiments were performed using cells from four different donors obtained from Lonza and n biological replicates as indicated in each figure where appropriate. Bar graphs are represented as mean ± SEM. Statistical analyses were performed using GraphPad Prism9, the threshold for significance level is set at P < 0.05. Detailed information on analyses is indicated in the figure legends.

## Supplementary Materials

See supplementary materials for gene expression data tables (Supplementary Tables S2-8) generated from RNAseq analysis of TGF-β1 and IL-13 treated microtissues.

## Acknowledgements

The authors would like to thank James Mainquist, Dr. Yingyao Zhou, Doug Quackenbush, and Dr. Jimmy Elliot for guidance in development of methodology and experimental design.

## Author Declarations

### Author contributions statement

Conceptualization: P.B., C.P., E.T., S.H., Methodology: P.B., C.P., C.C.S., P.C., Consumable Design and Fabrication: C.P., J.B, T.T., Data Generation and Analysis: P.B., C.P., J.B. C.C.S., P.C., B.L., O.R., J.W., S.W.B., Software Development: C.C.S. Writing Manuscript: P.B., C.P., S.H., E.T., Supervision: M.Q., S.H., E.T, Funding: Novartis Institute of Biomedical Research. All authors reviewed and approved the manuscript.

## Data availability

The RNAseq data reported in this paper will be uploaded (and the accession number provided) prior to paper acceptance. The data that supports the results in the paper are available in the article or supplementary information. Additional data for the figures are available from the corresponding author upon reasonable request.

---

**Supplemental Figure 1:**
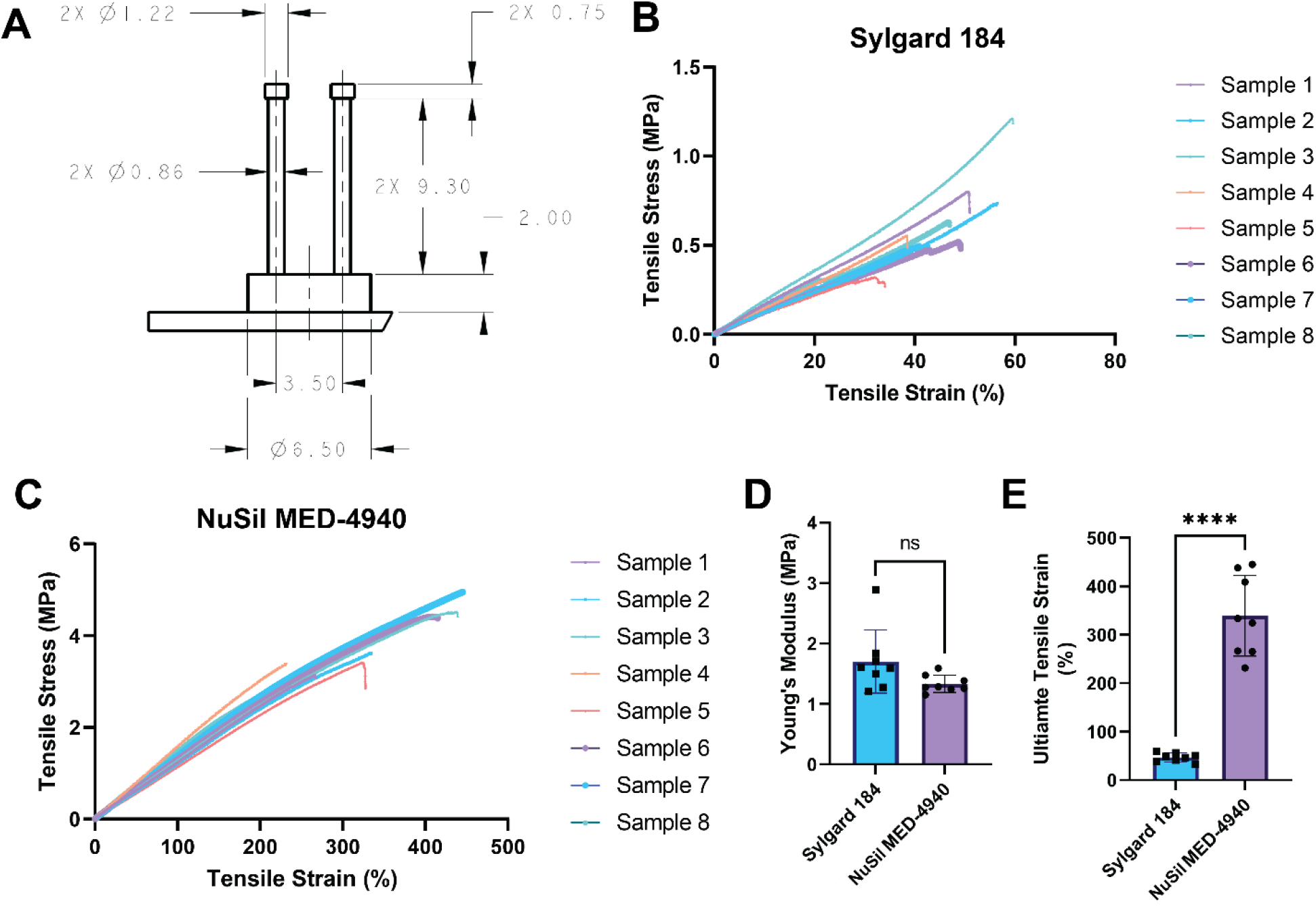
DEFLCT design and elastomer base enables high aspect ratio fabrication. **A)** Dimensional overview for DEFLCT cantilevers. All dimensions are shown in mm with global tolerances of ± 0.1mm for all callouts. **B)** Tensile stress-strain curves for rectangular samples of cast silicone elastomer Sylgard 184 and **C)** NuSil MED-4940. Samples were cut from several consumable fabrication batches and stretched until failure. **D)** Young’s modulus derived from the linear region of stress-strain curves for Sylgard 184 and NuSil elastomer. Data represented as mean ± S.T.D. **E)** Ultimate tensile stress of Sylgard 184 and NuSil elastomer determined by component failure. Data represented as mean ± S.T.D.

**Supplemental Figure 2:**
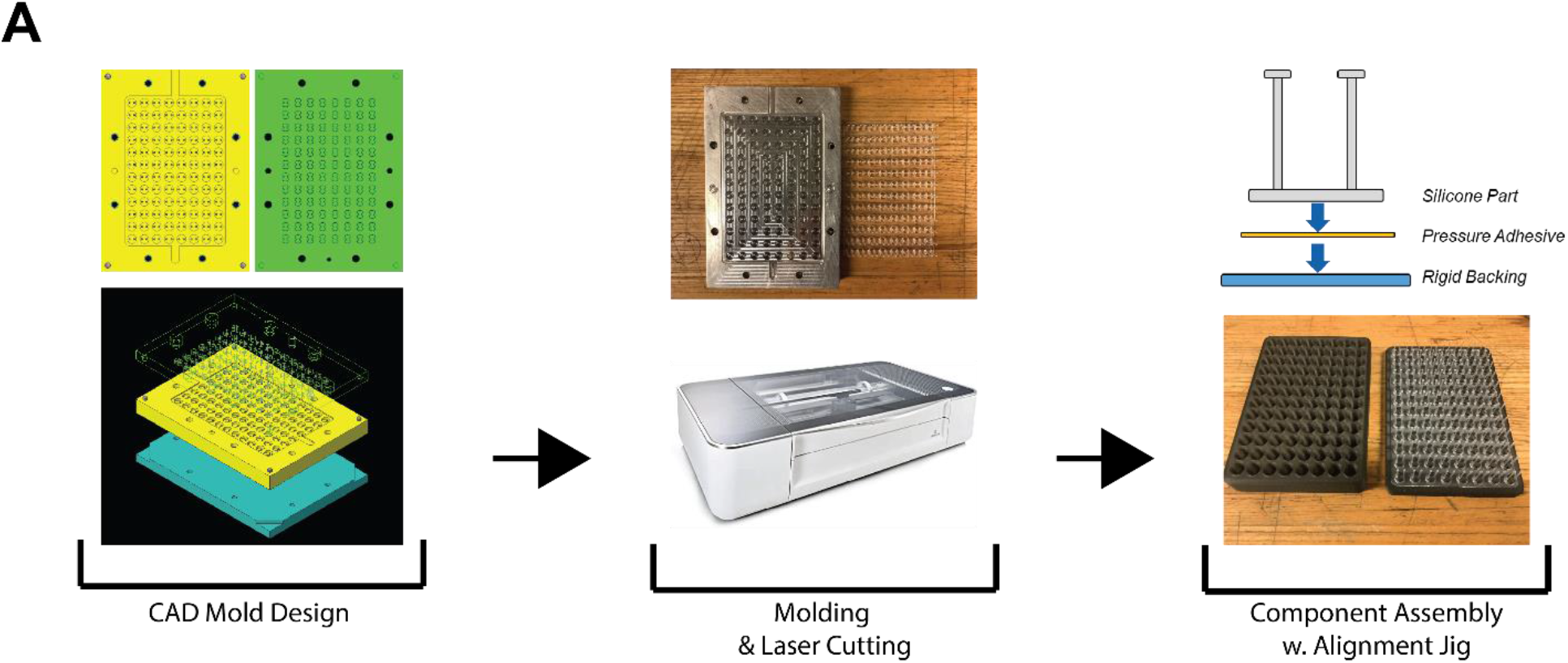
Graphical overview of DEFLCT component fabrication and assembly. **A)** Flowchart for component development. Aluminum mold sets for silicone arrays were designed in CAD software (top and bottom left) to match required dimensions. Mold sets were then machined and filled with liquid silicone resin under pressure and temperature to form a cured part within several minutes (top center). Simultaneously, laser cutting of the rigid acrylic backing for the elastomer’s structural support and precision cutting of pressure sensitive adhesive was carried out (bottom center). Finally, all components were assembled layer by layer (top right) using an in-house alignment tool (bottom right) to ensure through holes for pipetting are positioned correctly.

**Supplemental Figure 3:**
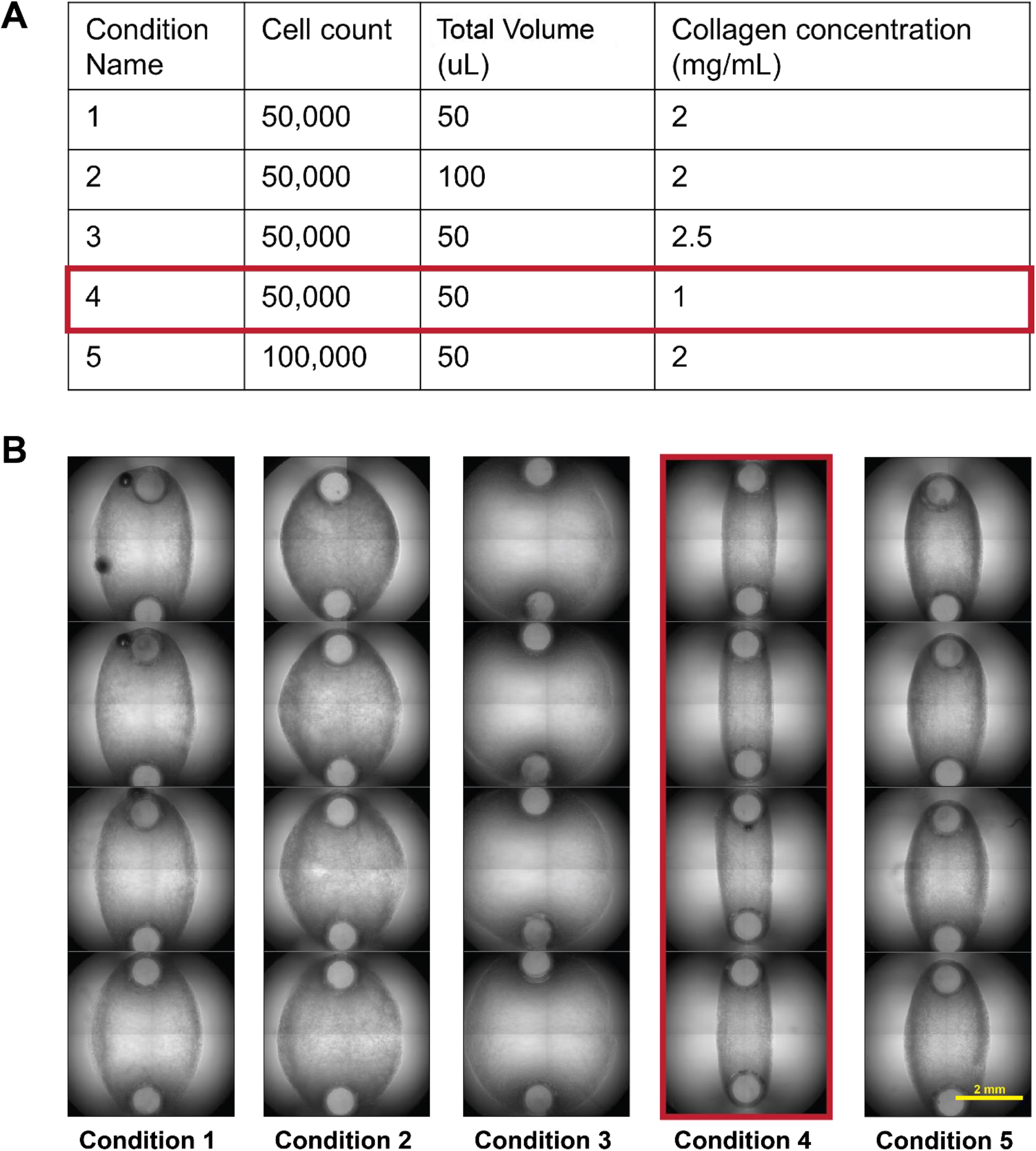
Smooth muscle tissue casting optimization on DEFLCT consumables. **A)** Tabular overview of experimental conditions for microtissue casting. Conditions vary in total tissue cell count, total volume of cell/ECM mixture, and baseline concentration of pre-tissue formation ECM in the seeding solution. **B)** Representative images of tissue formation given the above conditions. Optimal tissue formation was assessed by minimal total gel/tissue area after formation, clearly defined tissue borders, and early-stage deflection of cantilevers. Condition 4 (red) was selected for subsequent experiments. Scale bar = 2mm.

**Supplemental Figure 4:**
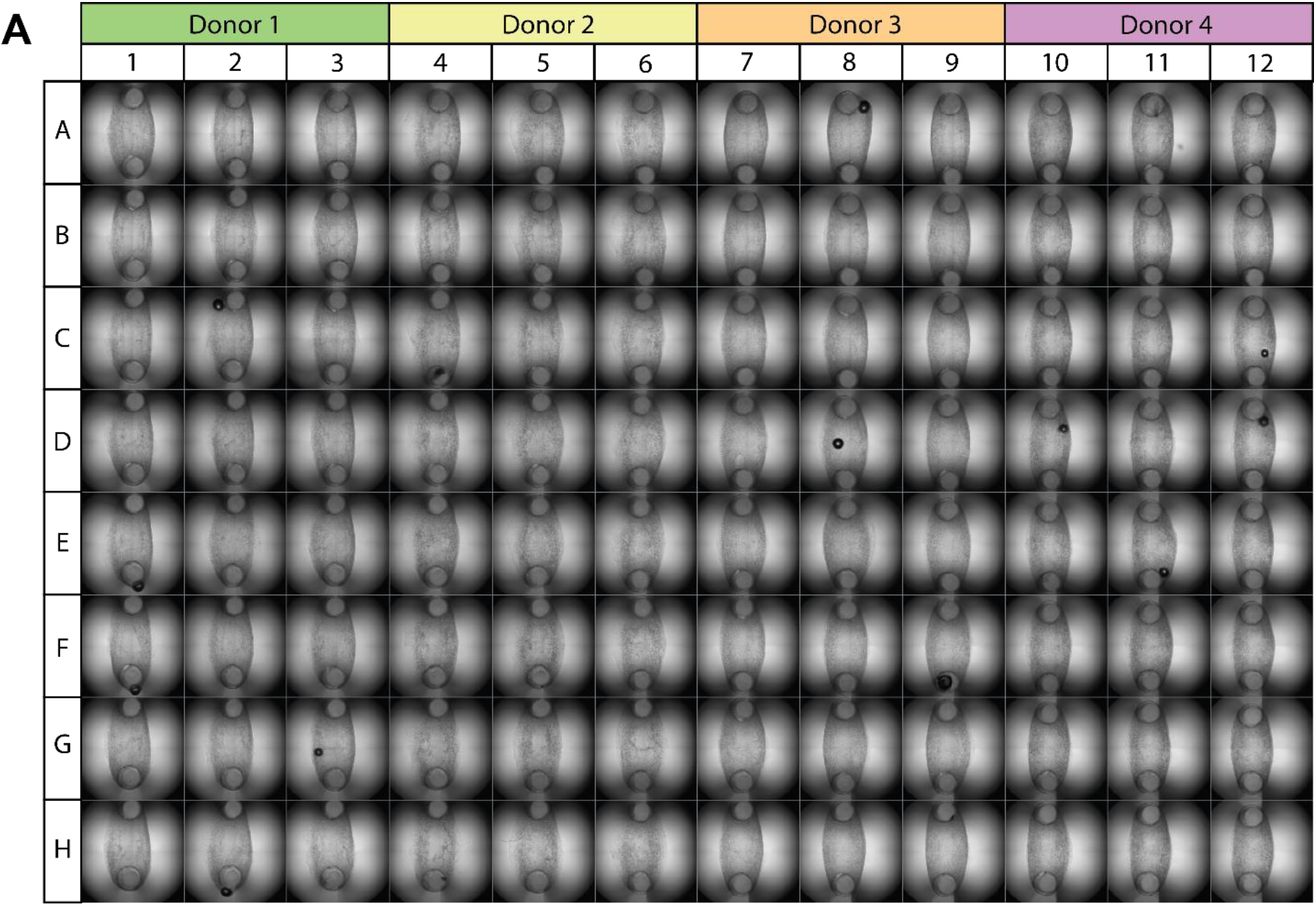
Full 96 well DEFLCT platforms can be easily seeded with microtissues. **A)** Maximum intensity projection brightfield images of a single DEFLCT consumable used in a standard 96 well format experiment. Plate map shows 4 separate hBSMC donors highlighted above their associated columns. Cantilever casting and tissue formation with a 100% success rate is typically observed following seeding. During treatment experiments all intra-plate replicate wells (organized here in groups of 3 columns) are averaged per cytokine treatment to produce single mean average datapoint representing one biological replicate.

**Supplemental Figure 5:**
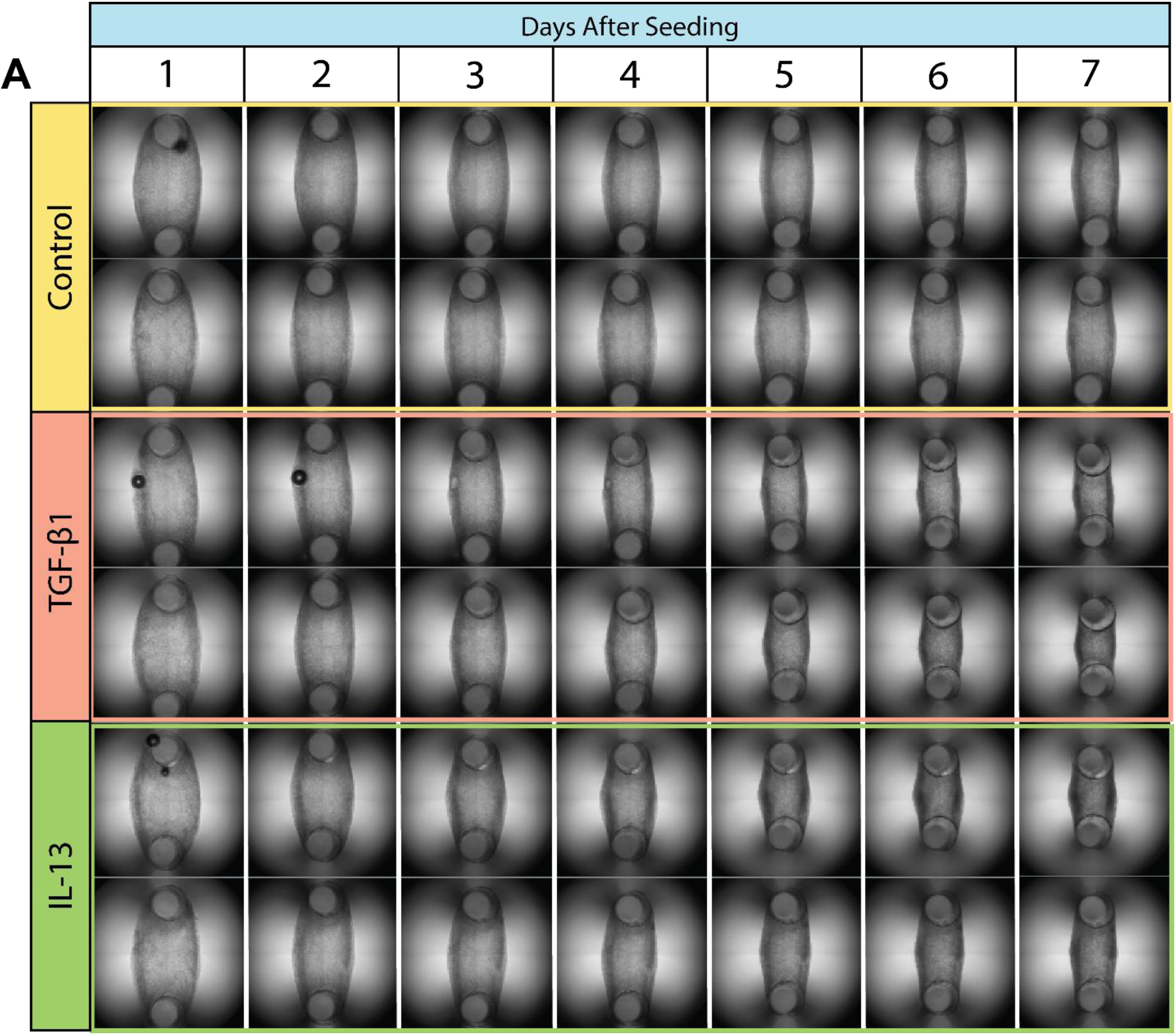
Tissue contraction with and without cytokines progresses over a 7-day period. **A)** Representative time course images taken daily over a 7-day period for control and contractility promoting cytokine treatment groups. Cantilever deflection across all groups begins to manifest between days 2 and 3, and the magnitude of contractility change per day appears to increase in cytokine treatment groups.

**Supplemental Figure 6:**
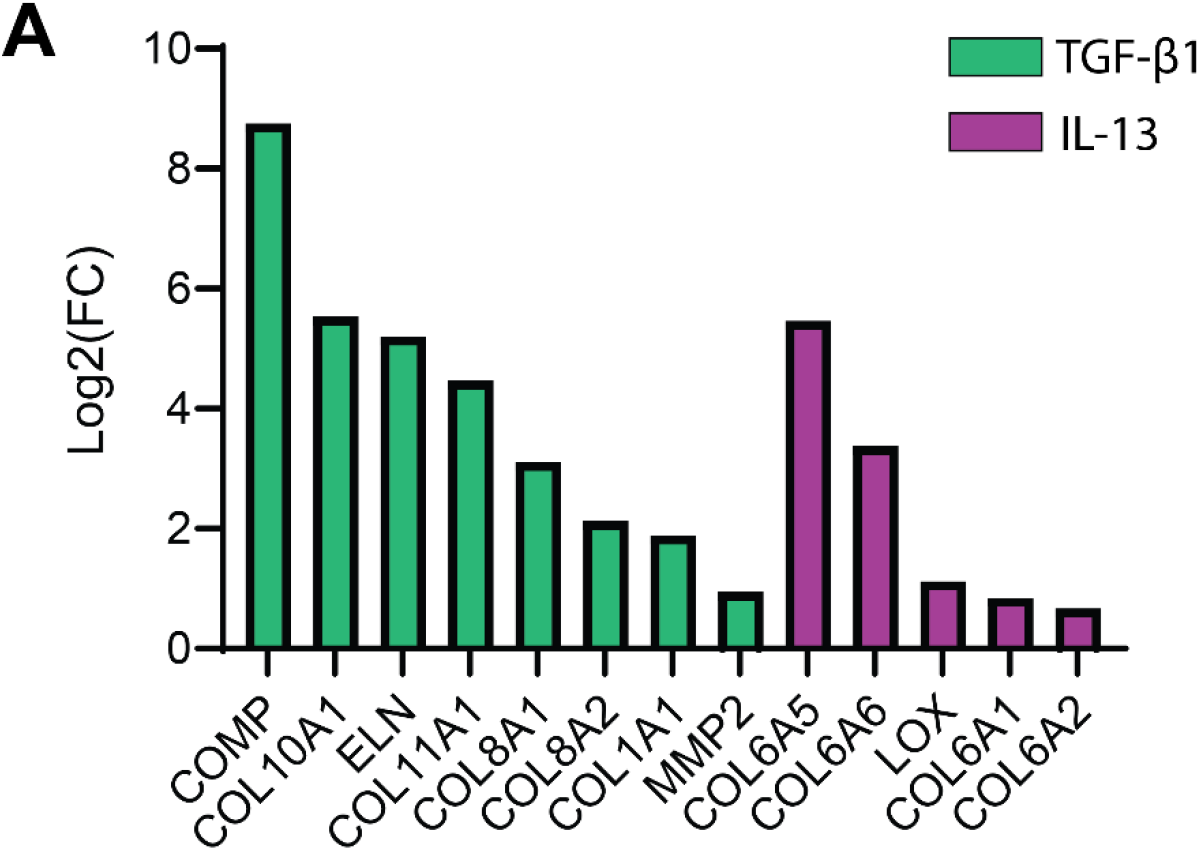
ECM remodeling increases after treatment with contractility inducing cytokines. **A)** Bar plot showing Log2(FC) readouts for top hit DEGs associated with extracellular matrix remodeling. DEGs for both TGF-β1 (green) and IL-13 (purple) are shown.

